# The ongoing invasion of the endogenous retrovirus *Kuruka* in natural *Drosophila melanogaster* populations

**DOI:** 10.1101/2025.10.01.679930

**Authors:** Riccardo Pianezza, Sarah Saadain, Matthew Beaumont, Sarah Signor, Robert Kofler

## Abstract

Transposable elements are mobile DNA sequences capable of proliferating within host genomes, occasionally capable of crossing species boundaries via horizontal transfer (HT). Here, we report the discovery and characterization of *Kuruka*, a newly invading endogenous retrovirus in natural *D. melanogaster* populations. *Kuruka* encodes an envelope protein and belongs to the *gypsy* /*gypsy* superfamily. Analysis of over 1000 *D. melanogaster* genomes revealed that *Kuruka* first appeared in sub-Saharan Africa in 2010. By 2017-2019 *Kuruka* had spread to Asia, Europe and America in 2017-2019, and is still actively invading Europe and Oceania as of 2021. Phylogenomic analyses suggest that *Kuruka* entered in *D. melanogaster* via a recent HT from an Afrotropical *Drosophila* species, most likely *D. erecta*. This is the first case of a recent HT from an Afrotropical donor species to *D. melanogaster*. In *D. erecta, Kuruka* has a single genomic insertion, which is located within *flamenco*, a master regulator of TE activity. The presence of an active host defense (piRNAs) suggests that *Kuruka* is silenced in *D. erecta*. Our findings establish *Kuruka* as a valuable model for studying the early stages of TE invasions and the dynamics of genome defense in real time.

## 1 Introduction

Transposable elements (TEs) are genomic sequences that hijack the host machinery to replicate themselves and increase in copy number within the genome. They are ubiquitous across the tree of life, having been detected in every eukaryotic genome studied to date [Wicker et al., 2007]. TEs fall into two major classes: retrotransposons, which use an RNA intermediate and a “copy-and-paste” mechanism to replicate, and DNA transposons, which rely on a “cut-and-paste” strategy without an RNA intermediate [Finnegan, 1989]. Retrotransposons can be further divided into long-terminal repeat (LTR) and non-LTR elements.

To control the propagation of TEs, eukaryotic hosts have evolved multiple molecular defense mechanisms to limit TE activity. In *Drosophila*, this defense is primarily mediated by the PIWI-interacting RNA (piRNA) pathway [Senti and Brennecke, 2010, Brennecke et al., 2007, Gunawardane et al., 2007]. piRNAs are small non-coding RNAs that recognize and silence TEs in a sequence-specific manner. They are typically generated from TE copies inserted within specialized genomic regions, known as piRNA clusters [Brennecke et al., 2007, Malone et al., 2009]. These clusters act as molecular ‘traps’: when a TE inserts into one, it triggers the production of piRNAs that can silence all members of the same TE family [Bergman et al., 2006, Brennecke et al., 2007, Zanni et al., 2013]. Distinct piRNA pathways operate in the germline and in the surrounding follicle (somatic) cells [Malone et al., 2009, Li et al., 2009a]. While both pathways rely on piRNAs, they are processed from different clusters that do not function identically. In somatic follicle cells, piRNAs are predominantly derived from a single master cluster, termed *flamenco*. In the germline, piRNAs are generated by many dispersed clusters [Pelisson et al., 1994, Prud’Homme et al., 1995, Malone et al., 2009, Brennecke et al., 2007, Yamanaka et al., 2014].

The evolutionary divergence of these two pathways reflects differences in TE expression: while most TEs are active in the germline, others have adapted to be expressed in somatic tissues. In fact, some TEs from the *gypsy/gypsy* superfamily contain an envelope gene, thought to be derived from DNA Baculoviruses [Terzian et al., 2001, Rohrmann and Karplus, 2001]. The acquisition of the envelope transformed these elements from intracellular replicating retrotransposons into infectious retroviruses [Malik et al., 2000]. These TEs, known as insect endogenous retroviruses (iERVs) or Errantiviruses, are typically active in the somatic tissue surrounding the germline [Senti et al., 2025]. Virus-like particles generated in the soma may infect the germline, and lead to novel TE insertions that will be transmitted to the next generation [Brasset et al., 2006]

Despite the presence of sophisticated host defense mechanisms, TEs often persist over evolutionary timescales. However, once silenced by the host, TEs will accumulate mutations and gradually degenerate. Some TEs manage to escape this fate through horizontal transfer (HT) into new species, where they can potentially re-establish activity in a naive genomic environment. Although the molecular mechanisms enabling HT of TEs are not fully understood, accumulating evidence shows that HT is widespread across eukaryotes [Zhang et al., 2020, Peccoud et al., 2017, Schaack et al., 2010, Bartolomé et al., 2009].

Until recently, HT was thought to occur on deep evolutionary timescales, with estimated rates of one invasion per species every several hundred thousand years [Bartolomé et al., 2009, Peccoud et al., 2017]. Indeed, only a few cases of recent HT have been documented [Bucheton et al., 1992, Periquet et al., 1989, Daniels et al., 1990, Schwarz et al., 2021, Scarpa et al., 2023, Pianezza et al., 2024a, Scarpa et al., 2025, Anxolabéh`ere et al., 1988]. The most striking examples are the recent invasion of 11 TEs in *D. melanogaster* during the last 200 years [Pianezza et al., 2025]. This rate of TE invasions is significantly higher than previously envisioned. Of these, three TEs were horizontally transferred from Neotropical *Drosophila* species, while the remaining eight are likely derived from *D. simulans*, the closest relative of *D. melanogaster* [Pianezza and Kofler, 2025]. Of the 11 recent invaders only *Tirant*, which spread in *D. melanogaster* populations around 1940, contains an envelope protein.

However, our ability to discover such recent invasions remains limited. Several of these invasions were discovered from phenotypic effects caused by TEs (hybrid dysgenesis) or a sudden increase in the number of copies of known TE families [Scarpa et al., 2023, Pianezza et al., 2024a, Anxolabéh`ere et al., 1988, Kidwell, 1983, Bucheton et al., 1992, Periquet et al., 1989]. We recently developed a novel approach that circumvents the need for prior knowledge of TE sequences. By identifying genomic regions present in assemblies of recently collected strains but absent from short-read data of older strains, it is possible to detect TE invasions that occurred between the two sampling points[Pianezza et al., 2025, 2024b]., such recent TE invasions may still be detectable through the extensive short-read data available for more recently collected strains [Chen et al., 2024]. However this approach exclusively identifies invasions present in the investigated assemblies. Since suitable long-read assemblies are available only for strains collected before 2018, it is possible that very recent invasions were missed [Rech et al., 2022, Pianezza et al., 2025]. Nevertheless, the extensive short-read data from more recently collected strains could still capture such recent events [Chen et al., 2024]. Using a novel pipeline to detect TE invasions from short-read sequencing data, we discovered a new iERV currently spreading in *D. melanogaster* populations, which we have named *Kuruka*. Phylogenetic and genomic analyses suggest that *Kuruka* entered the *D. melanogaster* genome via a recent HT from *D. erecta*. With high resolution spatio-temporal data, we reconstructed the detailed timing of *Kuruka*’s spread across global *D. melanogaster* populations. This work establishes *Kuruka* as a new model for studying both the dynamics of iERV invasions and the real-time genomic responses of host populations.

## 2 Results

### 2.1 The discovery of *Kuruka* based on short-read data

To test whether novel TE invasions were captured by short-read data from recently sampled *D. melanogaster* strains we developed a novel pipeline. The idea is that a very recent TE invasion should lead to TE sequences being present in short-read data of recently collected strains but absent in the assemblies of older strains. Our pipeline thus aims to identify TE sequences that are absent in a reference genome. A similar strategy had previously been used to identify DNA viruses in *Drosophila* sequencing data [Wallace et al., 2021]. Briefly, short-reads are mapped to the reference genome, and unmapped or poorly mapped reads are extracted. Reads from known contaminants (e.g., *Homo sapiens, Wolbachia*) are filtered out, and the remaining reads are assembled *de novo*. The assembled contigs are aligned to a TE protein database and high-confidence matches are retained. Final candidates are manually curated based on metrics such as coverage depth and sequence length. A schematic overview of the pipeline is shown in Fig. S1.

We applied this pipeline to several recently collected *D. melanogaster* samples [Chen et al., 2024]. Our pipeline successfully recovered previously documented invasions of TEs such as *P* -element, *Spoink, McLE*, *Souslik*, and *Transib1*. This confirms that our pipeline may detect invasions of TEs not present in a reference assembly. In addition to these known TEs, we detected a previously unannotated ∼9kb sequence that is frequently present in samples collected from China between 2017 and 2020.

To generate a high-quality consensus of this unknown element, we sequenced wild-caught flies from North Dakota (USA) in 2023 on a MinION long-read platform. The raw reads contained 12 sequences matching this element. We generate a consensus sequence of this novel element based on a multiple sequence alignment of the 12 sequences (Fig. S2).

A BLASTn search did not reveal a significant match with any known *D. melanogaster* TE family [Rech et al., 2022, Quesneville et al., 2005, Hubley et al., 2016], except for weak similarity (∼68% identity over ∼20% of the sequence) to both *Gypsy5* and *Tirant*. We refer to this novel element as *Kuruka*, derived from the Swahili word for “jump”.

### 2.2 Kuruka is an endogenous retrovirus of the *Gypsy* /*Gypsy* superfamily

*Kuruka* is 8833 bp long and features two perfectly identical LTRs of 514 bp. Based on ORF prediction (ORFinder) and BLASTp analysis, it encodes a *gag* protein, a *pol* polyprotein, and an envelope (*env)* protein (Fig. 1a).

**Figure 1:**
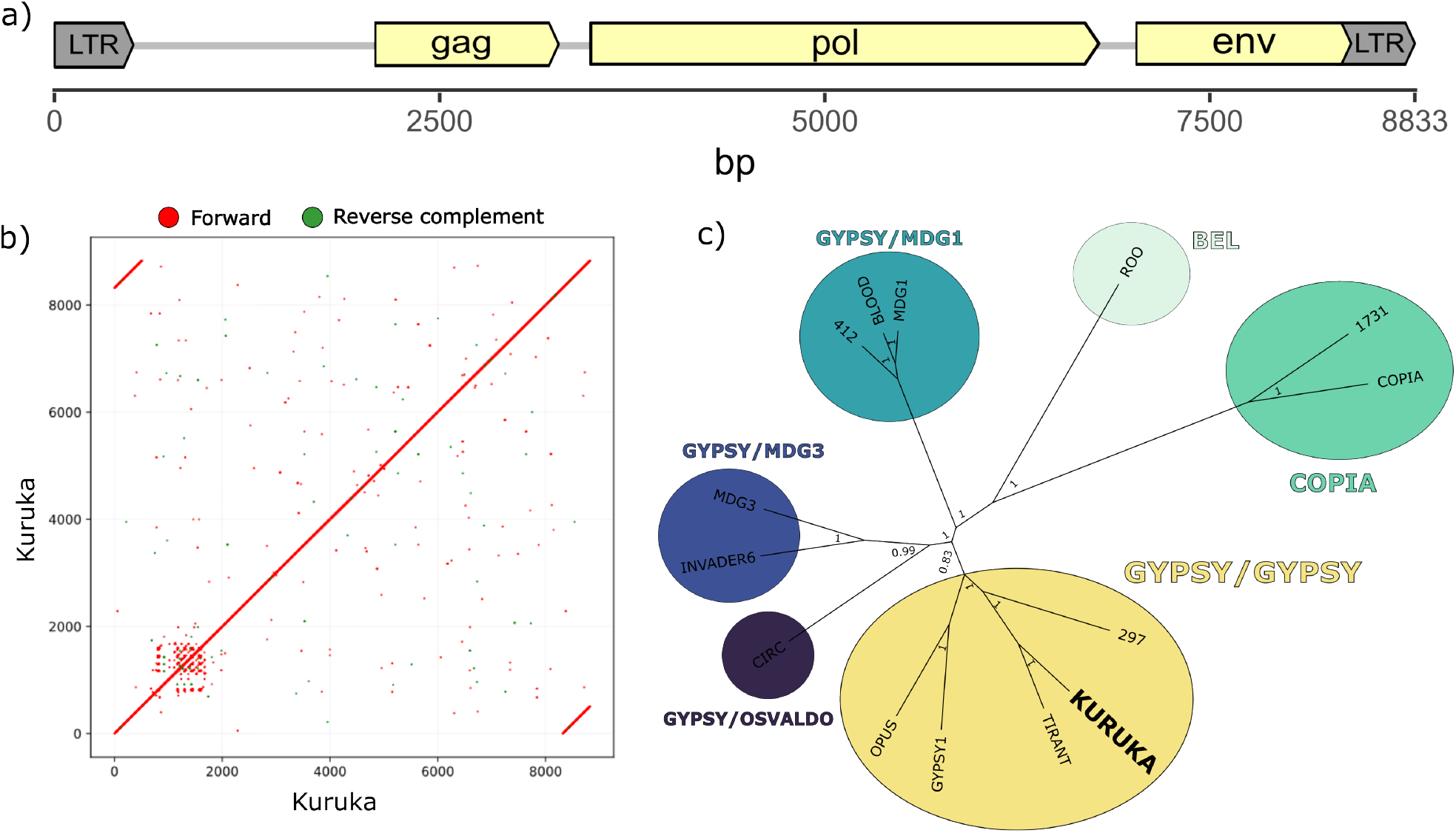
Structural and phylogenetic characterization of *Kuruka*. a) Features of the 8833 bp *Kuruka* consensus sequence, showing two LTRs and protein-coding domains for *gag, pol*, and *env*. b) Dot plot of the *Kuruka* sequence, highlighting the two LTRs and a low-complexity region within the first 2 kb. c) Bayesian phylogenetic tree of the reverse transcriptase domains from known *D. melanogaster* TEs and *Kuruka. Kuruka* clusters within the *Gypsy* /*Gypsy* superfamily.

A dot plot of *Kuruka* against itself revealed two LTRs, but also a ∼ 800bp repetitive region located within the first 2kb (Fig. 1b). This low-complexity region lies within the non-coding portion of the element. A 313 bp portion of this repetitive region is present in some insertions and absent in others (Fig. S2), representing a structural variant among insertions of the TE family.

To determine the classification of *Kuruka*, we constructed a Bayesian phylogenetic tree based on the reverse transcriptase domains of known *D. melanogaster* TEs (Fig. 1c). The resulting tree places *Kuruka* within the *Gypsy* /*Gypsy* superfamily, close to *Tirant*. This classification is consistent with the presence of an *env* gene in the sequence of *Kuruka* [Kapitonov and Jurka, 2003]. *Similar to other Gypsy* /*Gypsy* transposons, *Kuruka* may thus be active in the somatic follicle cells around the ovary.

### 2.3 Timing and geographic spread of *Kuruka* in *D. melanogaster* populations

To reveal the spatio-temporal spread of *Kuruka* in *D. melanogaster* populations, we analyzed 1072 publicly available short read datasets [Grenier et al., 2015, Schwarz et al., 2021, Lange et al., 2021, Rech et al., 2022, Kapun et al., 2021, *Shpak et al., 2023, Pool et al., 2012, Chen et al., 2024, Nunez et al., 2025]. The sequenced flies were were collected from* ∼ *1815 (museum collections) through 2021 from populations across the globe. This genomic time-series data enables us to explore the spread of Kuruka* at a high-resolution.

We used DeviaTE [Weilguny and Kofler, 2019] to estimate the copy number of *Kuruka* in each sample based on coverage depth. DeviaTE estimates the copy number of a TE by normalizing the coverage of the TE by the coverage of single copy genes. We considered *Kuruka* to be present in a sample if 90% of its sequence had a normalized coverage > 1.

Our time-series data revealed that *Kuruka* was completely absent from all samples collected before 2010 (Fig. 2a). It first appeared in 2010 but was not detected again until 2017, when it reached its maximum estimated copy number (∼ 50). In subsequent years, *Kuruka* was consistently present in some samples while remaining absent in others (Fig. 2b).

**Figure 2:**
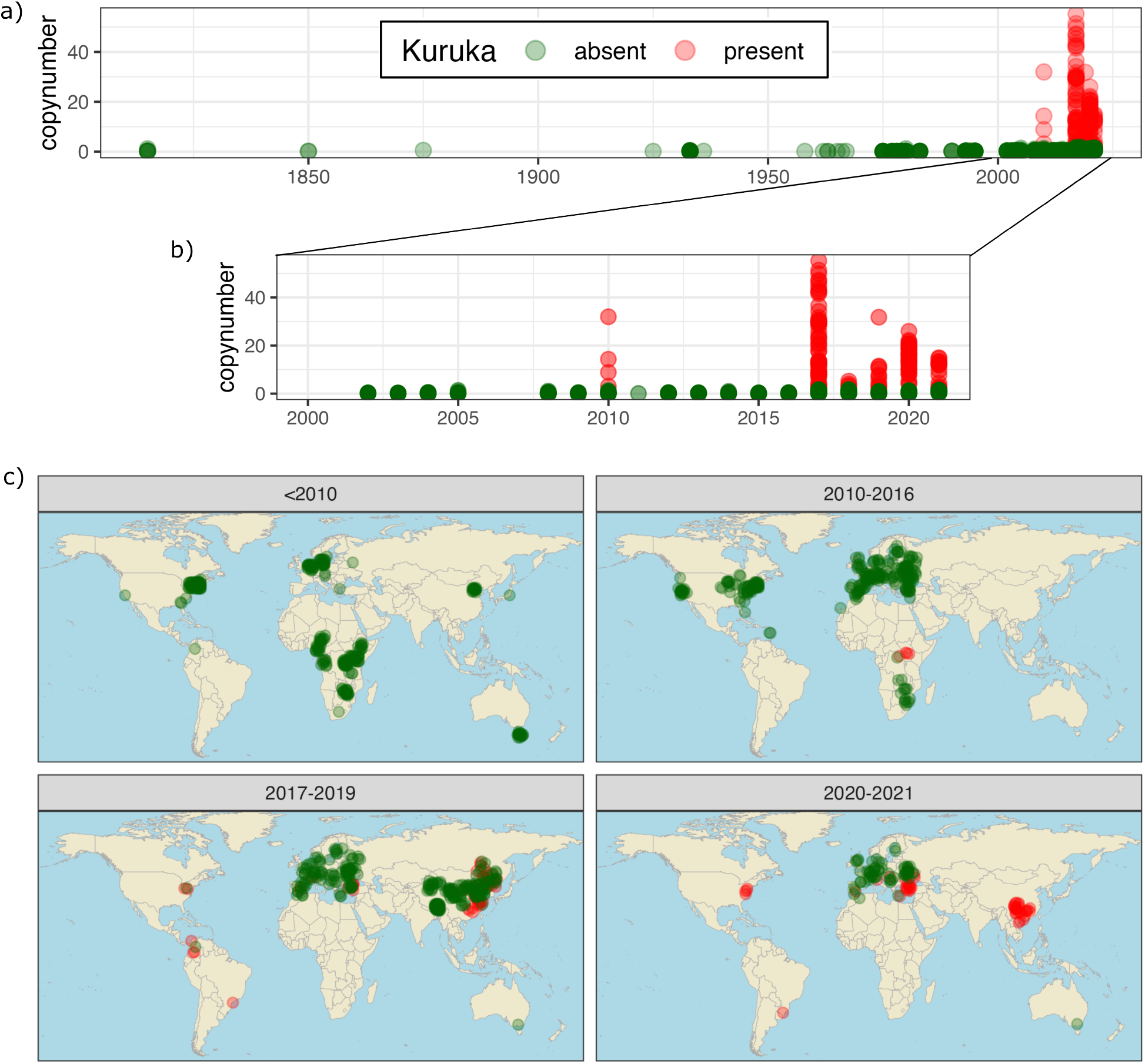
Temporal and geographic spread of *Kuruka* in *D. melanogaster* populations. a) Timeline of the *Kuruka* invasion, based on short-read data. The presence (red) and absence (green) of *Kuruka* was inferred from the normalized coverage. *Kuruka* is absent in all samples prior to 2010, and present in some (but not all) samples thereafter. b) Zoomed-in view of the timeline from 2000 to 2018. From 2017 onward, *Kuruka* is found each year in at least a few samples. c) The invasion of *Kuruka* in natural *D. melanogaster* populations. *Kuruka* is first detected in samples from Africa in 2010. Around 2017–2019, *Kuruka* is also found in samples from Asia and the Americas, and sporadically in Europe. By 2020–2021 *Kuruka* is found in the Americas, East Asia and multiple samples from Europe but not in Australia.

To better understand this peculiar temporal distribution, we examined the geographic origin of the samples (Fig. 2c). *Kuruka* first appeared in Africa in 2010, but was not detected elsewhere until 2017. Between 2017 and 2019, *Kuruka* began to spread into East Asia, the Americas, and a few samples from Europe. By 2020–2021, all samples from the Americas and East Asia carried *Kuruka*, while it remained absent from Australian samples and roughly half of the European samples. Our data suggest that in 2021 the *Kuruka* invasion was still ongoing in natural *D. melanogaster* populations.

As additional time-points, our newly collected and sequenced samples from 2023 in North America (ND1-4) had *Kuruka* insertions.

We also sequenced two iso-female lines of *D. melanogaster* collected in 2025 in North Dakota and Washington, D.C. using ONT, and they both carried multiple *Kuruka* insertions. We identified at minimum 16 full-length insertions of *Kuruka* across all major chromosome arms (per haplotype).

We sought to determine whether *flamenco* insertions of *Kuruka* are present in this recently collected *D. melanogaster*, which would be a sign that the TE is now silenced. In the strain collected in Washington, D.C we found a *Kuruka* insertion in *flamenco*, however this was absent in the strain from North Dakota. This could mean that *Kuruka* is only partially silenced in American populations by 2025 or that other insertions triggered the silencing of *Kuruka*.

### 2.4 Origins of *Kuruka*

Because *Kuruka* first appears in *D. melanogaster* populations from Africa, we hypothesized that the donor species responsible for the HT initiating the *Kuruka* invasion may also have an African origin. To identify the donor species, we searched for the *Kuruka* sequence in over 1500 arthropod genomes, including 49 *D. melanogaster* assemblies, 305 other drosophilids, and 1211 non-drosophilid arthropods.

Since *Kuruka* has never been reported in *D. melanogaster*, we did not expect to find it in any genome of this species. Only a fragment resembling *Kuruka* was detected in 35 out of 49 of the *D. melanogaster* assemblies. This fragment is 4 kb long, 11% diverged from *Kuruka* and contains no LTRs (Fig. S3). The insertion site of this sequence is identical in each of the 35 assemblies, suggesting an ancient origin.

Among the non-drosophilid arthropods, there is no relevant match for *Kuruka* (Fig. S4). The situation is drastically different in drosophilids: several species show strong matches with *Kuruka*, both in terms of alignment length and sequence identity. The best matches were found in *D. erecta* (8658bp with 98.85% sequence identity) and *D. bocqueti* (7497bp with 98.96% sequence identity). Interestingly, all species having insertions strongly resembling *Kuruka* are Afrotropical *Drosophila* species of the melanogaster group (Fig. 3A).

**Figure 3:**
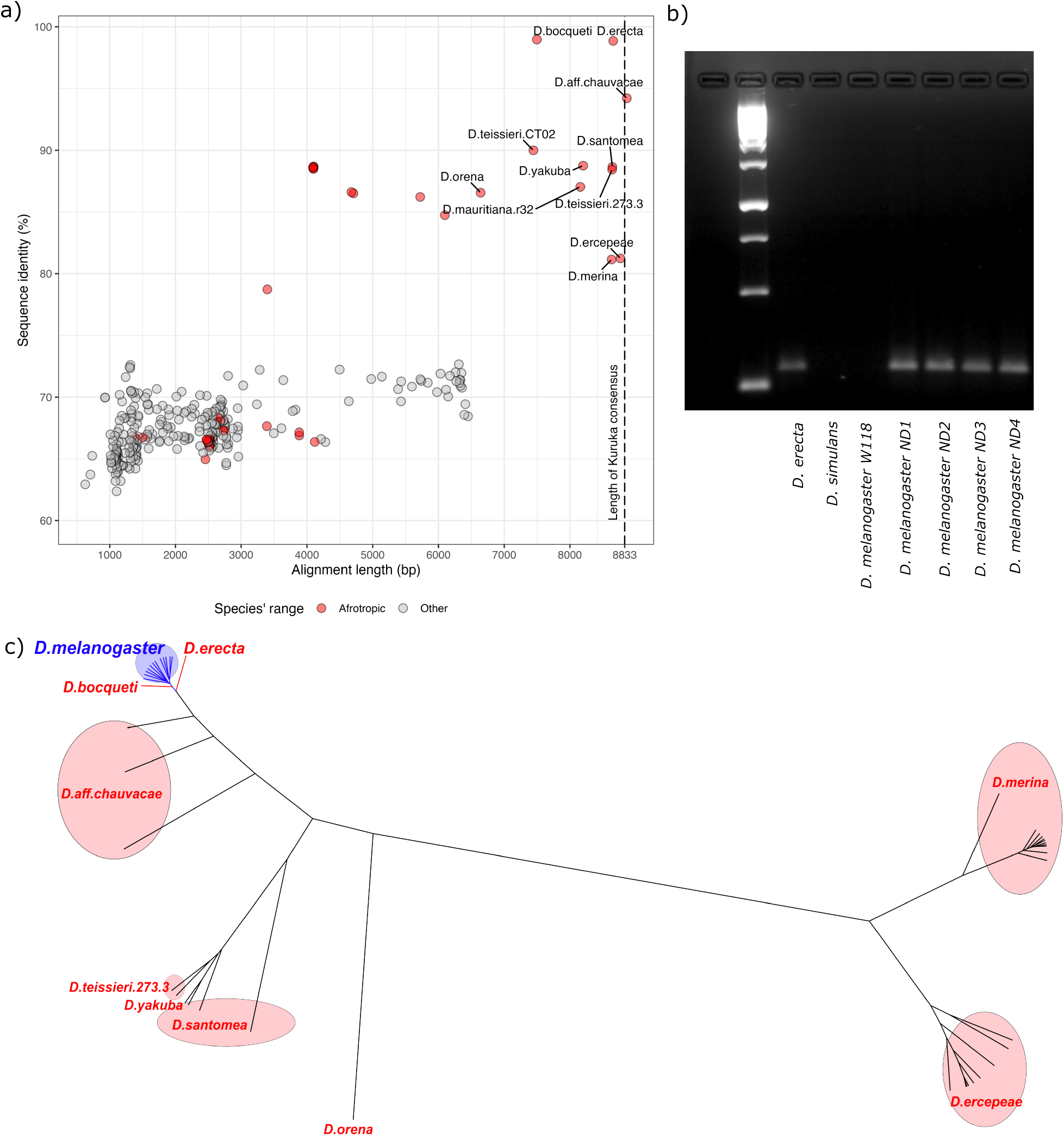
Origins of *Kuruka*. a) Sequences resembling *Kuruka* in 334 drosophilid assemblies. The graph shows alignment length and sequence similarity between *Kuruka* and the best match in each species. Species are colored by their ancestral biogeographic realm. Insertions with the highest sequence similarity to *Kuruka* were found in *D. erecta* and *D. bocqueti*, where the match is longer in *D. erecta*. b) PCR confirms that *Kuruka* is present in *D. erecta* and a *D. melanogaster* strain sampled in 2024, yet absent from *D. simulans* and an older *D. melanogaster* strain (w^118^). c) Bayesian phylogenetic tree of full-length *Kuruka* insertions across multiple *Drosophila* species. The 12 insertions identified in our ONT data (*D. melanogaster* strain collected 2023 in North Dakota) form a monophyletic clade, consistent with a recent expansion from a single introduction. Insertions from *D. erecta* and *D. bocqueti* are most closely related to the *D. melanogaster* insertions.

To validate the presence–absence pattern of *Kuruka* in selected *Drosophila* species, we performed PCR (Fig. 3B). PCR confirmed our previous results, with *Kuruka* being found in *D. erecta*, as well as in *D. melanogaster* samples collected in 2023 (ND1-4), while it was absent from an older *D. melanogaster* strain (*w*^*118*^ and *D. simulans* [Qiu et al., 2017]. *This patchy taxonomic distribution, with Kuruka* being present in *D. melanogaster* and the distantly related *D. erecta* but absent in the closely related *D. simulans*, is a classic signature of HT [Bartolomé et al., 2009, Loreto et al., 2008].

To further investigate the potential donor species, we constructed a Bayesian phylogenetic tree with all full length-insertions of *Kuruka* across all species (Fig. 3C). The tree also includes the 12 *Kuruka* insertions in *D. melanogaster* identified in our long-read data. These 12 insertions form a monophyletic group and cluster closely together, confirming their low sequence divergence and supporting the hypothesis of a recent invasion. Located very close to the *D. melanogaster* clade were insertions from *D. erecta* and *D. bocqueti*. To our surprise, only a single insertion of *Kuruka* was found in each of these two species. The Bayesian tree solely considers the matching nucleotide sequences of the alignment and ignores indels. Given that a substantially larger portion of *Kuruka* aligns with *D. erecta* (8658 bp) than with *D. bocqueti* (7497 bp), we argue that *D. erecta* is the most likely donor.

In summary, *Kuruka* is found exclusively in *D. melanogaster* and in African *Drosophila* species within the *melanogaster* group. The *Kuruka* invasion was most likely triggered by HT from an African *Drosophila* species, with *D. erecta* being the most probable donor.

### 2.5 The single *Kuruka* insertion in *D. erecta* likely generates piRNAs in somatic cells

The likely donor species, *D. erecta*, contains a single *Kuruka* insertion. Although somewhat unexpected, it is not impossible that a species with just a single TE insertion could trigger a novel invasion in *D. melanogaster*. We wanted to investigate the *Kuruka* insertion in *D. erecta* in further detail. First, we confirmed that only a single insertion of *Kuruka* is present in *D. erecta*. To do so, we re-sequenced and assembled the *D. erecta* strain 14021-0224.01 that was used to generate the reference genome. The new assembly contains fewer contigs than the reference, while showing higher N50 and N90, (see Table S1), indicating improved completeness and continuity. This newly generated assembly of *D. erecta* still contains only a single *Kuruka* insertion (Fig. 4A).

**Figure 4:**
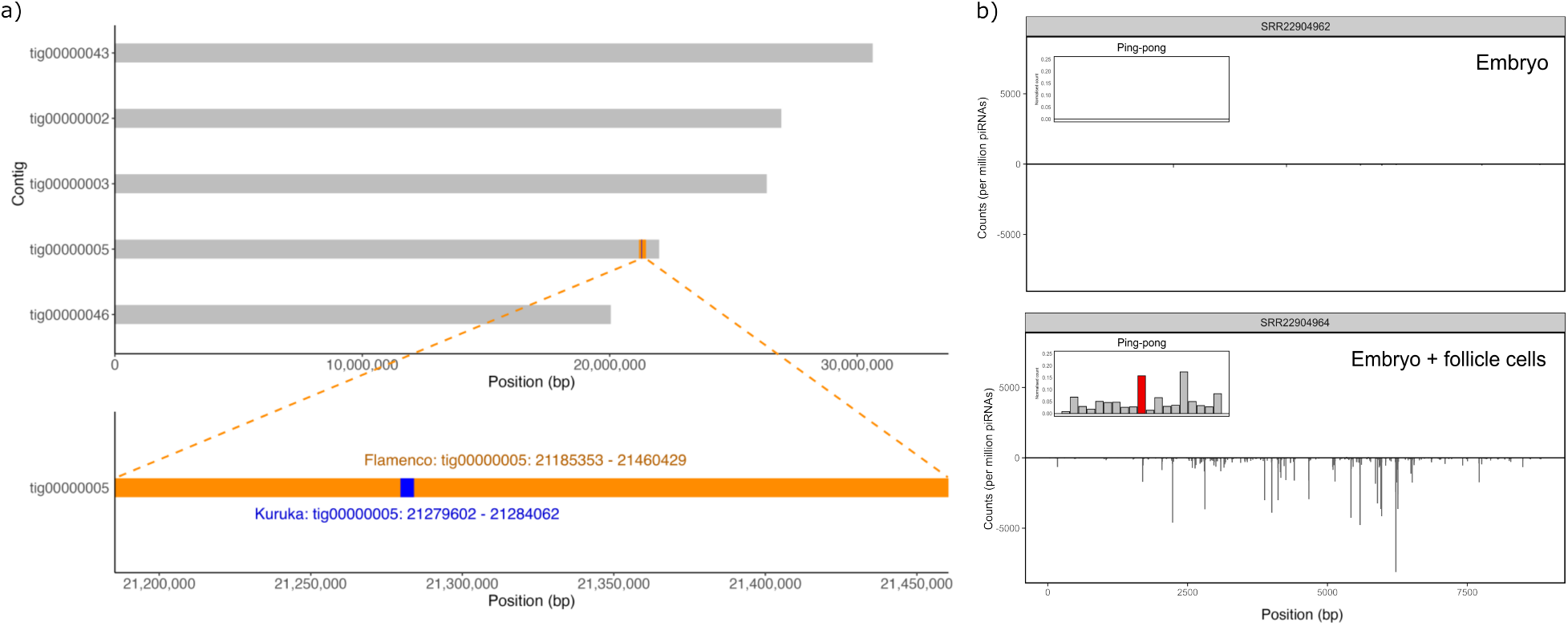
The single *Kuruka* insertion in *D. erecta* is located in *flamenco* and generates piRNAs. (a) A single insertion of *Kuruka* is found in our novel *D. erecta* genome assembly, consistent with the reference genome. The insertion is located within the *flamenco* locus. (b) We found antisense *Kuruka* piRNAs in the ovary (germline + soma) but not in the embryo (germline) of *D. erecta* flies. A weak ping-pong signature in ovarian piRNAs suggests that the germline pathway may contribute to silencing of *Kuruka*. Our data suggest that *Kuruka* is largely silenced by the somatic piRNA pathway in *D. erecta*.

The *Kuruka* insertion in *D. erecta* is located within *flamenco* (Fig. 4A, see Materials and Methods), a somatic piRNA cluster. This is in agreement with *Kuruka* likely being active in the soma (as suggested from the presence of an envelope protein and its classification in the *Gypsy* /*Gypsy* superfamily). The insertion in *flamenco* further suggests that *Kuruka* is silenced by the somatic piRNA pathway in *D. erecta*.

To further investigate this, we analyzed publicly available small RNA data extracted from different *D. erecta* tissues [Selvaraju et al., 2024]. We mapped piRNAs to the *Kuruka* consensus sequence, along with other TE families present in *D. erecta* as controls. In samples derived from ovaries, which contain piRNAs from both the germline and the somatic pathway, many piRNAs aligned to *Kuruka* (Fig. 4B, S5). By contrast, in the embryo, which only contains piRNAs from the germline, very few small RNAs map to *Kuruka* (Fig. 4B). This suggests that *Kuruka* is largely silenced by the somatic piRNA pathway. As a control, other TEs are targeted by piRNAs in both the ovary and the embryo (Fig. S6). In ovaries we found a weak ping-pong signature for *Kuruka*, which is a hallmark of the germline piRNA pathway (Fig. 4B). It is unclear if the germline piRNA pathway contributes to silencing of *Kuruka*.

To summarize, the potential donor of *Kuruka, D. erecta*, has a single *Kuruka* insertion in *flamenco* and *Kuruka* is likely silenced by the piRNA pathway in *D. erecta*.

## 3 Discussion

Our work reveals the ongoing invasion of a new iERV, which we named *Kuruka*, in natural populations of *D. melanogaster*. Phylogenetic and genomic evidence strongly supports a HT event likely originating from *D. erecta*, an Afrotropical species of the *melanogaster* group. This represents the 12th documented TE invasion in *D. melanogaster* over the past 200 years.

Although our pipeline for finding novel TEs in short read data identified the *Kuruka* invasion and recovered previously known cases (e.g., *P* -element, *Spoink*), it is possible that it may have missed other invading TEs. Detecting recent TE invasions using short reads is challenging due to their limited length and the potential inclusion of exogenous sequences (e.g., bacterial, viral), which complicates the identification of novel TEs, especially those present at low copy number. In contrast, our tool GenomeDelta [Pianezza et al., 2024b] enables more accurate detection of newly invading TE families, but it requires high-quality long-read data of recently collected strains, which are scarce. Although we consider it unlikely, we can therefore not rule out that additional TEs may have spread in *D. melanogaster* very recently.

In this work we precisely reconstructed the spatiotemporal spread of *Kuruka* in natural *D. melanogaster* populations. This level of resolution was only possible thanks to the global sampling efforts of projects like DrosEU and DrosRTEC [Kapun et al., 2021, Nunez et al., 2025], which have generated and made available hundreds of natural population datasets over time. Sustained initiatives of this kind are critical for identifying and characterizing additional ongoing TE invasions in real time.

By 2021 *Kuruka* was still absent in Oceania and some European populations, while it spread in the Americas and East Asia. This suggests that the *Kuruka* invasion was still ongoing by 2021. Also the absence of *Kuruka* insertions in *flamenco* in one of the two strains sampled in 2025 in North America suggests that the *Kuruka* invasion is not yet fully silenced by the piRNA pathway. However, we cannot rule out that the host defense has been triggered by alternative mechanism, such as antisense insertions of *Kuruka* in 3’ UTRs of host genes [Rafanel et al., 2025].

With the likely donor being *D. erecta* (or another Afrotropical *Drosophila* species), *Kuruka* is the first HT event in *D. melanogaster* in the last 200 years that does not originate from a species of the *D. simulans* complex or the *D. willistoni* group. While phylogenetic studies have suggested many ancient HT of TEs from other Afrotropical drosophilid species into *D. melanogaster* [Carareto, 2011, *Bartolomé et al., 2009, Modolo et al., 2014, Pianezza and Kofler, 2025], Kuruka* is the first documented case that has occurred during the last 200 years. This further shows that the high-rate of recent TE invasions in *D. melanogaster* and some related Drosophilidae is not limited to a very few donor species. The habitat expansion of *D. melanogaster* into the Americas during the past 100-200 years, which was suggested as cause for several of the previous invasions (including invasions where the cosmopolitan *D. simulans* acted as donor, e.g. due to potent vectors of HT with habitats in the Americas), can not explain all recent invasions [Daniels et al., 1990, Scarpa et al., 2025, Pianezza et al., 2024a, 2025].

Remarkably, we identified only a single *Kuruka* insertion in the putative donor species, *D. erecta*, where the TE is likely silenced by the somatic piRNA pathway. A single *Kuruka* insertion is also found in the potential alternative donor species, *D. bocqueti*, but we do not have any information about the piRNAs in this species. However, assuming that *D. erecta* is the donor, this raises important questions about the dynamics of *Kuruka* in this species, and the mechanisms that enabled its successful invasion of *D. melanogaster*. Two main hypotheses could explain the presence of only one insertion in the donor species. First, it is possible that *Kuruka* invaded these genomes long ago, and that all insertions except the single insertion in *flamenco* were gradually lost through genetic drift or negative selection. Alternatively, *Kuruka* may have inserted early on into a piRNA-producing locus which would have triggered its immediate silencing and prevented further spread. It is feasible that some somatic TEs have a strong insertion bias into *flamenco*. For example, *Shellder*, a TE that spread in *D. simulans* within the last 50 years, has multiple independent insertions in *flamenco*. Moreover, many independent *Tirant* insertions in *flamenco* were found in different *D. melanogaster* strains [Rafanel et al., 2025]. *Notably, Tirant* is closely related to *Kuruka* and also invaded *D. melanogaster* fairly recently (∼ 1940 [Schwarz et al., 2021]). A strong insertion bias of somatic TEs into *flamenco* could halt the invasions of somatic TEs at their earliest stages. If *D. erecta* is the donor of the HT, our results would imply that a silenced TE, with just a single copy in a genome, may still be horizontally transmitted and trigger a novel invasion in a naive host. This is not impossible, given that the host defense (e.g. piRNAs) is likely not transmitted along with the TE during a HT. However, we cannot exclude the possibility that the actual donor lineage was a different species, or an unsampled population of *D. erecta*.

*Kuruka* is only the second TE, after *Transib1*, for which the spatio-temporal dynamics of invasion have been precisely reconstructed [Pianezza et al., 2025]. Interestingly, *Kuruka* seems to spread much slower in natural populations than *Transib1*. Despite first appearing in 2010, *Kuruka* has not spread to the whole European population as of 2021. By contrast, the DNA transposon *Transib1*, spread globally within just two years from its initial appearance in 2014 [Pianezza et al., 2025]. Why is *Kuruka* spreading slower than *Transib1* ? Interestingly, *Tirant*, a somatic TE closely related to *Kuruka* that invaded *D. melanogaster* around 1940 is still absent in some strains sampled > 2000, i.e. 60 years after the start of the invasion [Rech et al., 2022, Pianezza et al., 2025, Rafanel et al., 2025]). This suggests that *Tirant* may also be spreading slowly in natural *D. melanogaster* populations. *Kuruka* and *Tirant* are the only iERVs encoding envelope proteins among the 12 recent invaders. Their unique life cycle, requiring somatic expression, production of virus-like particles, and transfer to the germline, may impose biological constraints that limit transposition rates and slow down global spread. We can speculate that somatic TEs (i.e. *Kuruka* and *Tirant*) spread more slowly than TEs that are active in the germline (e.g. *Transib1*).

Together, our findings show that the high rate of recent TE invasions in *D. melanogaster* is still ongoing and that we can expect further TE invasions in the next years. Our work positions *Kuruka* as a powerful new model to study the invasion dynamics of iERVs and the interplay between TEs and the host silencing system in real time.

## 4 Materials and Methods

### 4.1 Detection of *Kuruka* in short-read datasets

Short-read datasets from various *D. melanogaster* strains collected in China in 2017 and 2020 [Chen et al., 2024] were mapped to the reference genome (GCA 000001215.4) using bwa mem (v0.7.17-r1188) [Li and Durbin, 2009] with default parameters. Unmapped reads were extracted using samtools (v1.13) [Li et al., 2009b], and reads with more than 5% sequence divergence were extracted using a custom Python3 script (badmapped-finder.py). The reads were then filtered to remove potential contaminants by aligning with a custom database containing genomes of common laboratory contaminants (e.g., *Homo sapiens, Wolbachia*) using bwa mem. The remaining reads were assembled *de novo* using SPAdes (v3.13.1) [Prjibelski et al., 2020]. The assembled contigs were queried against a transposable element protein database [Novak, 2023] (v3.1, metazoa) using BLASTx. Only contigs with high confidence TE-related hits were retained for further curation (sequence identity > 80%, alignment length > 250bp).

### 4.2 Long-read sequencing of *D. melanogaster*

New iso-female lines of *D. melanogaster* were collected in 2023 and 2025 (col. Tim Greives and David Wright, respectively). For the flies collected in North Dakota in 2023 (*ND1-3*), three of these lines were pooled and used for long read sequencing at North Dakota State University on a MinION platform (Oxford Nanopore Technologies, Oxford, GB) with base calling using Dorado (1.0, ARM64). Two additional iso-female lines from 2025 were sequenced separately (*DW7* and *DC1*) after being collected in Fargo and Washington D.C. They were sequenced and base called using the aforementioned pipeline. These reads were assembled using hifiasm [Cheng et al., 2021, 2025].

### 4.3 Consensus sequence reconstruction and structural annotation

To detect *Kuruka* insertions in the raw long-reads, we used RepeatMasker (v4.1.2-p1) [Smit et al., 2013-2015] with options -no is -s -nolow and a custom TE library containing only the contig with the *Kuruka* sequence assembled from the short-reads. Only insertions longer than 80% of the query sequence were retained and aligned using MAFFT (v7.526) [Katoh et al., 2002].

To get a consensus sequence we used a Python3 script (MSA2consensus.py), which applies a majority-rule approach to assign a nucleotide at each position in the multiple sequence alignment.

Protein domains encoded in the resulting consensus sequence were identified using ORFfinder, and subsequently aligned against the NCBI protein database using BLASTx to identify similar proteins [Wheeler et al., 2003]. The dot plot was generated using VectorBuilder [VectorBuilder Inc., 2025].

### 4.4 Estimating copy numbers using short-reads datasets

We estimated the abundance of *Kuruka* in 1072 publicly available *D. melanogaster* short-read datasets [Grenier et al., 2015, Schwarz et al., 2021, Lange et al., 2021, Rech et al., 2022, Kapun et al., 2021, Shpak et al., 2023, Pool et al., 2012, Chen et al., 2024, Nunez et al., 2025]. We first aligned the reads to a database consisting of the consensus sequences of *Kuruka* and three single copy genes (BUSCO IDs: 29at7147, 591at7147, 898at7147) with bwa bwasw (v0.7.17-r1188) [Li and Durbin, 2009]. To measure copy numbers, we used our tool DeviaTE [Weilguny and Kofler, 2019], which estimates the copy number of a TE by normalizing the coverage of the TE by the coverage of the single copy genes.

### 4.5 Long-read sequencing and genome assembly of *D. erecta*

Ovaries from ∼ 300 *D. erecta* (strain 14021-0224.01) were collected in ice-cold 1×PBS and high-molecular-weight DNA was extracted using the Monarch Genomic DNA Purification Kit (T3060, New England Biolabs). Library preparation and sequencing were performed by the Vienna BioCenter Core Facilities using the Oxford Nanopore Ligation Sequencing Kit V14 (SQK-LSK114) and a PromethION flow cell. Basecalling was done using Guppy (v6.5.7+ca6d6af).

Raw reads were filtered to 100× genome coverage and assembled with Canu v2.2 [Koren et al., 2017]. The assembly was polished through three rounds of Minimap2 (v2.30) [Li, 2018] and Racon (v1.5.0) [Vaser et al., 2017], then refined with Pilon (v1.24) [Walker et al., 2014] using Illumina reads from *D. erecta* populations published by Selvaraju et al. [2024].

To locate *flamenco* in the newly generated *D. erecta* assembly, we relied on the coordinates of *flamenco* in the *D. erecta* reference genome [van Lopik et al., 2023]. We retrieved the sequence of *flamenco* and mapped it to our assembly using bwa mem (v0.7.17-r1188) [Li, 2013].

### 4.6 Identification of *Kuruka* in genome assemblies

We investigated the presence of *Kuruka* in 49 *D. melanogaster*, 305 drosophilids, and other 1211 arthropod genome assemblies. For an overview of all analysed assemblies, see Supplementary File S1. We identified insertions in these assemblies using RepeatMasker (v4.1.2-p1; -no-is -s -nolow) [Smit et al., 2013-2015] and a custom library containing the consensus sequence of *Kuruka*.

For each drosophilid species in our dataset, we manually assigned its ancestral biogeographic realm based on Markow and O’Grady [2005] and if no information was available for a given species, the portal GBIF [GBIF, 2024].

### 4.7 Bayesian phylogenetic analysis

To infer the phylogenetic classification of *Kuruka*, we identified the RT regions in the consensus sequences of known *D. melanogaster* TEs and *Kuruka* using BLASTx against the NCBI protein database. We performed a multiple sequence alignment with MAFFT (v7.526) [Katoh et al., 2002] and a Bayesian tree was generated with the BEAST package (v2.7.5) [Bouckaert et al., 2019].

To generate a phylogenetic tree for the *Kuruka* insertions in the different *Drosophila* species, we used the RepeatMasker output and bedtools (v2.30.0) [Quinlan and Hall, 2010] to extract the sequences of *Kuruka* including 3kb flanking regions. Then, we used the LTRharvest tool from GenomeTools (v1.6.5) [Gremme et al., 2013] to select only insertions with two LTRs, removing any flanking region and fragmented insertions. The trees were built with MAFFT and BEAST [Katoh et al., 2002, Bouckaert et al., 2019].

### 4.8 PCR amplification of *Kuruka* insertions

PCR primers were designed based on *Kuruka* sequence in *D. erecta*. The primers are as follows: (F CCC-CTCCTGCTTGTTTACGT, R TGGTACGGTCAACTTCCAGC Invitrogen). The presence of *Kuruka* was screened in DNA from *D. erecta* (14021-0224.01), *D. simulans* (*SZ129* [Signor et al., 2018]), *D. melanogaster* from prior to the *Kuruka* invasion (*w*^*118*^), and four *D. melanogaster* strains collected in Fargo, North Dakota in the Fall of 2023. *D. erecta* functions as a positive control, while *D. simulans* and *D. melanogaster w*^*118*^ are negative controls.

### 4.9 Estimation of piRNA production

The abundance of piRNAs, the distribution of piRNAs within the *P* -element and the ping-pong signature were computed using previously described Python scripts [Kofler et al., 2018, Selvaraju et al., 2024]. *D. erecta* TEs used as controls were identified by annotating TEs *de novo* in the *D. erecta* reference genome using EarlGrey (v4.4.5; option -c yes) [Baril et al., 2024].

## Acknowledgments

R.P. would like to thank Pauline King’ori for her linguistic assistance in naming the TE, and Almor`o Scarpa for the precious scientific discussions.

## Author contributions

R.P. conceived the project and discovered the invasion of *Kuruka*. R.P., S.Sa., M.B. and S.Si. analysed the data. S.Sa. extracted the *D. erecta* genomic DNA and generated the genome assembly. S.Si. collected *D. melanogaster* strains, performed ONT sequencing and performed PCR. R.P. and R.K. wrote the manuscript. S.Sa., M.B. and S.Si. contributed to writing. R.K. supervised the project.

## Data Availability

The newly assembled *D. erecta* genome and raw reads for *D. melanogaster* lines ND1-4 are available at NCBI under accession number xxx. The analysis performed in this work have been documented with RMarkdown and have been made publicly available, together with the resulting figures and the library of *D. erecta* TE families consensus sequences generated with EarlGrey, at GitHub (https://github.com/rpianezza/Kuruka).

