## Supplementary Material for "The ongoing invasion of the endogenous retrovirus *Kuruka* in natural *Drosophila melanogaster* populations"

### Supplementary figures and tables

#### Supplementary figures

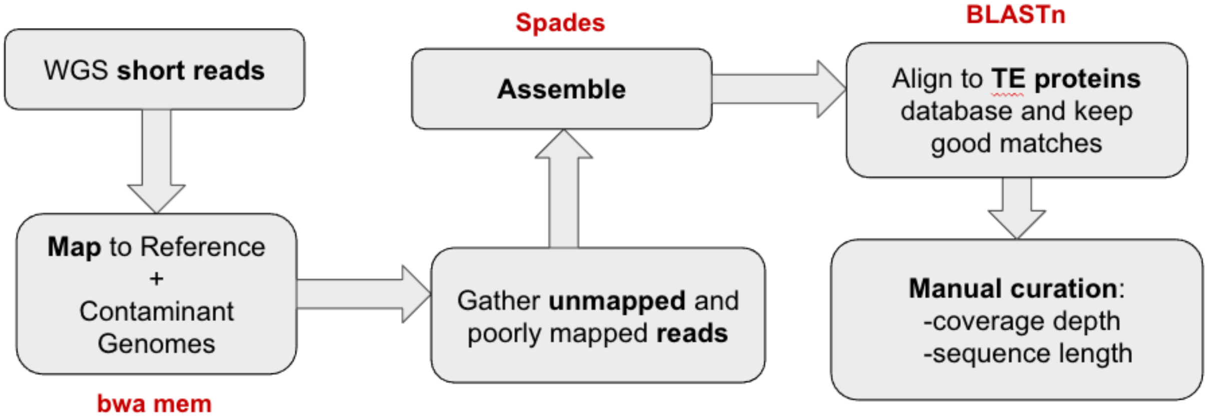


Figure 1: Schematic of the pipeline used to identify potential non-reference TE families, like *Kuruka*.

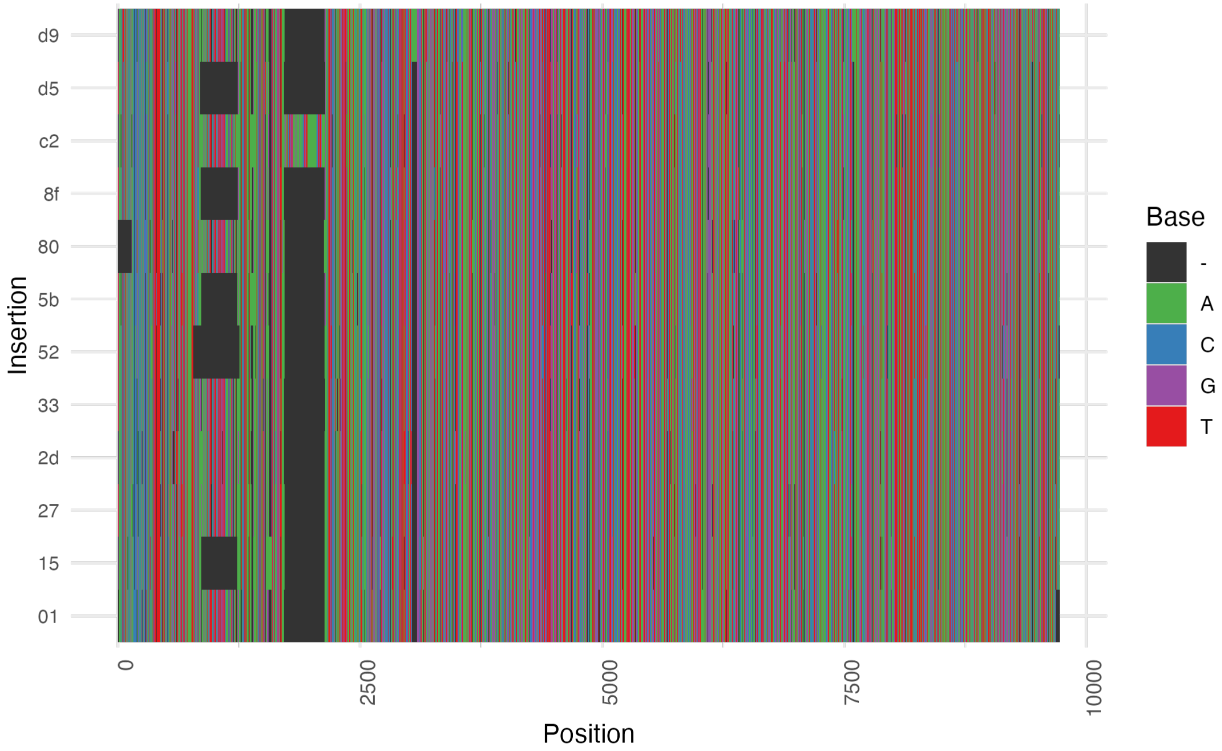


Figure 2: Multiple sequence alignment of the 12 *Kuruka* insertions found in the long reads sequenced from the *D. melanogaster* samples collected in 2023 in North Dakota. A common structural variant is evident around position 1000.

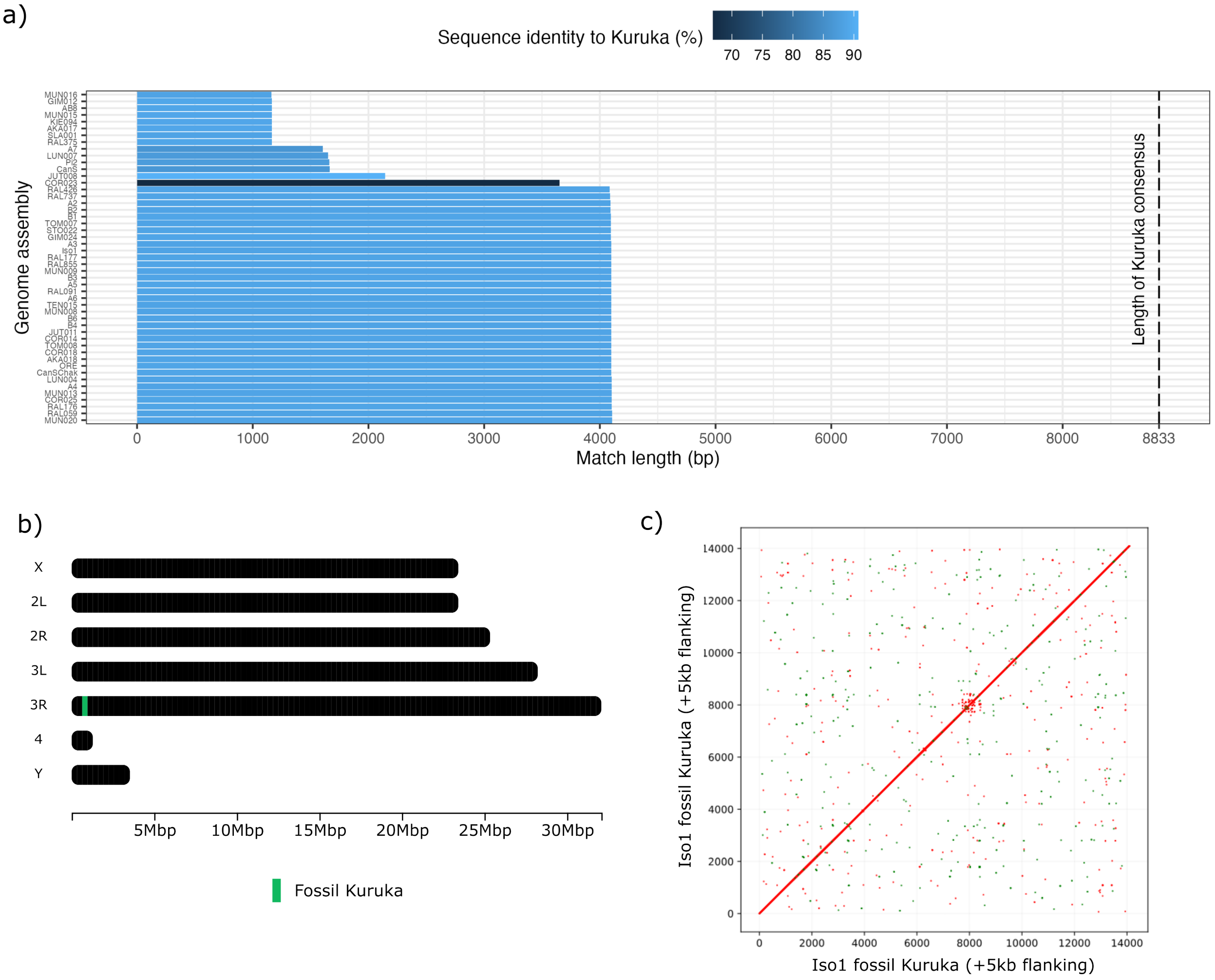


Figure 3: a) Best *Kuruka* hits identified in 49 *D. melanogaster* genome assemblies, including the reference genome (Iso-1). A ∼4 kb sequence sharing ∼88% identity with *Kuruka* is found in 35 genomes; remaining assemblies contain only lower-similarity fragments. This sequence is referred to as the "fossil" *Kuruka*. b) The fossil *Kuruka* is consistently located at the distal end of chromosome arm 3R, inserted at the same genomic position in all 35 genomes where it is detected. c) No LTRs are detected in the fossil *Kuruka* sequence, even after examining 5 kb of flanking regions on both sides.

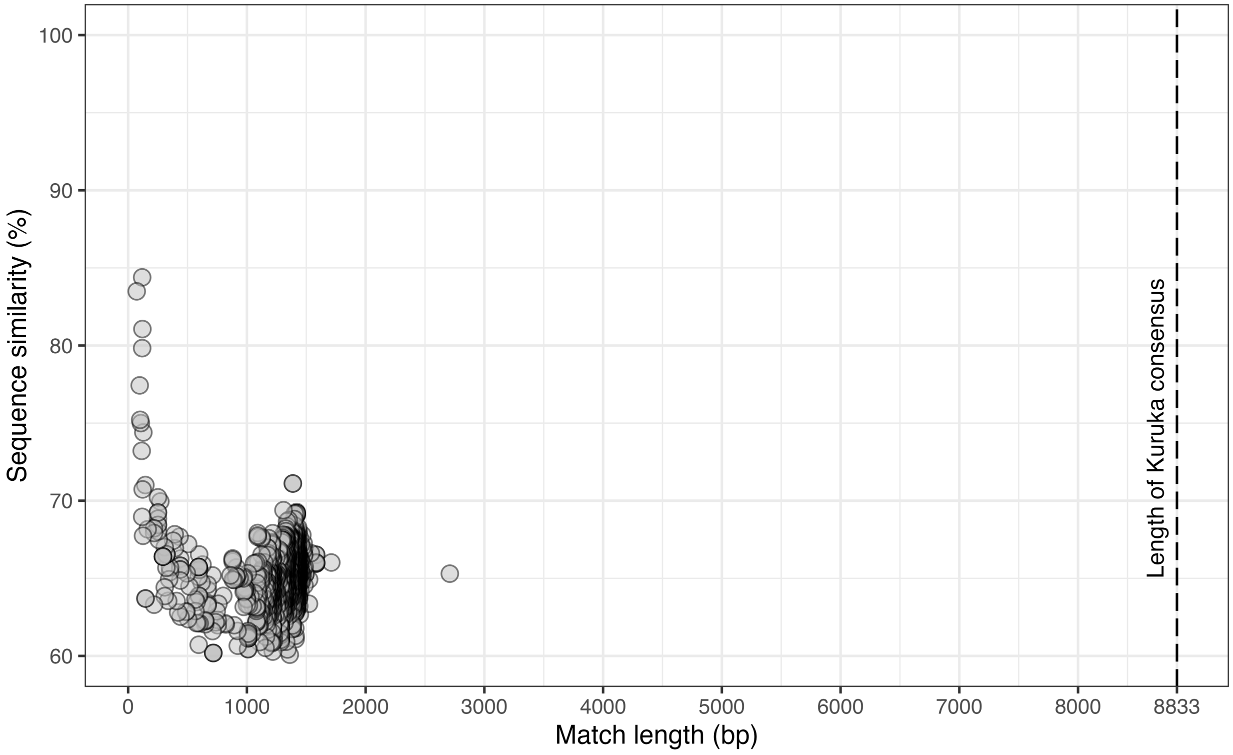


Figure 4: Best *Kuruka* hits identified in 1211 non-drosophilids arthropods genome assemblies. No relevant match to any of the genomes can be highlighted.

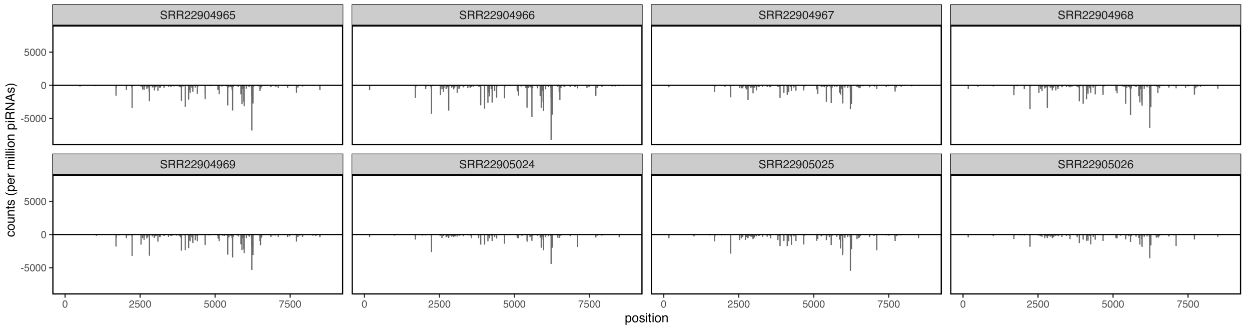


Figure 5: piRNA distribution over the *Kuruka* sequence for samples not shown in the main figure. Note that all of these samples contain both germline and somatic cells.

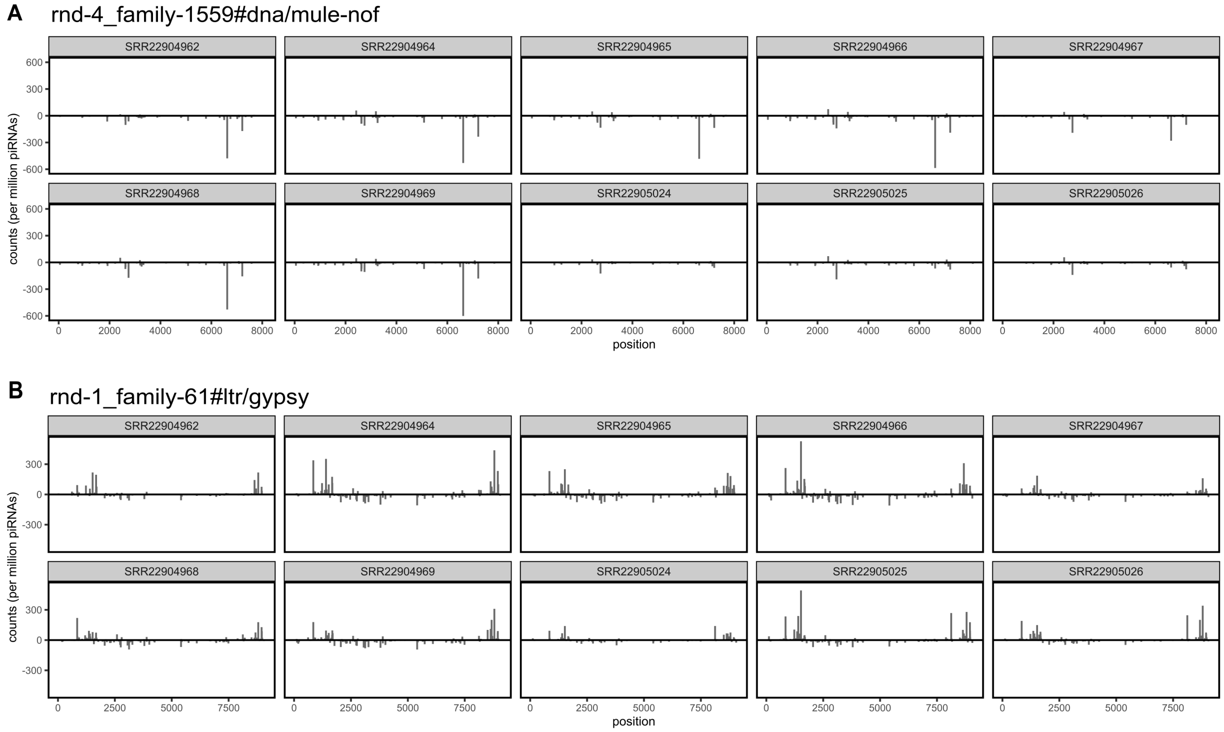


Figure 6: piRNA distribution on the sequences of two *D. erecta* TE families, used as positive controls. Both the TEs have piRNAs consistently mapped in all of the samples, validating the difference between embryo and soma highlighted for *Kuruka*.

#### Supplementary tables

Table 1: The newly assembled *D. erecta* genome has less contigs but higher N50 and N90 compared to the reference genome. BUSCO found the same percentage of orthologous genes in both the assemblies.

| | **GCA_003286155.2 (reference genome)** | **GCA_xxx (newly assembled genome)** |
|---|---|---|
| Contigs | 94 | 49 |
| N50 (bp) | 28426653 | 30625941 |
| N90 (bp) | 778323 | 1614603 |
| Complete BUSCOs (%) | 99.7 | 99.7 |

### References
